# The uptake of attractant and flavour enhanced baits by foxes (*Vulpes vulpes*) in rural England

**DOI:** 10.1101/147249

**Authors:** G.C. Smith, J.A. Woods

## Abstract

Over a 15 month period the removal rates of different attractant and flavour enhanced baits, by foxes, was investigated at permanent and single use sites. All baits were composed of mechanically recovered chicken meat (MRM) and were treated with one of four additives (an attractant or flavour); untreated baits were used as experimental controls. The addition of synthetic ferment egg (SFE) increased bait removal compared with untreated and valeric acid (VA) treated bait. However, the addition of beef flavour or honey flavour to bait did not increase bait removal rates compared to untreated bait.

There was limited evidence of learning (by foxes and other species responsible for removing the bait) at one of the permanent study sites but no evidence of such behaviour at the other site. There was no evidence of consistent seasonal differences in bait removal rates.

It was concluded that the use of attractants may increase bait removal rate in areas where bait removal rate is low; but where bait removal rate is high (in most rural areas of Britain) the addition of attractants or flavours offers little advantage.

## INTRODUCTION

For wildlife rabies control to be effective by oral vaccination it is necessary to attract a high proportion of the reservoir species to the bait (Anderson 1986; Smith & Harris 1989; Smith & Cheeseman 2002) and the proportion required for successful control is related to host density (Smith & Wilkinson 2003). The red fox (*Vulpes vulpes*) has been the main wildlife rabies reservoir in Europe, and thus the main concern for the UK, particularly as density has increased in urban areas (Wilkinson & Smith 2001; Scott *et al.* 2014), and the other potential hosts (raccoon dogs *Nyctereutes procyonoides* and raccoons *Procyon lotor*) are not present. Field trials have shown that should an outbreak of wildlife rabies occur in urban areas of Britain where fox density is high, bait uptake may be low enough to hamper disease eradication (Smith & Harris 1991; Trewhella *et al.* 1991; Smith & Woods 2007). Bait uptake rates in urban areas of Australia (>80%: Marks & Bloomfield 1999), rural Europe (>60%: Schneider 1985; Pastoret & Brochier 1998) and rural Britain (>60%; unpublished data) is higher and may be sufficient to quickly control an outbreak. Consequently to ensure the eradication of an outbreak of wildlife rabies in the UK, methods of increasing uptake of bait by foxes were investigated. Since many urban fox populations have a food surplus (Harris & Woollard 1988) a potential method to increase bait uptake by foxes would be to make the bait more attractive than other food sources. Saunders and Harris (2000) investigated a number of attractants and flavours against captive foxes and concluded that two attractants (synthetic fermented egg and valeric acid) and two flavours (beef and honey) merited further investigation as additives to improve bait uptake by foxes.

This initial work, performed in the early 1990s, looked for ways to increase bait uptake rates to improve the UK rabies contingency plans (Harris, Smith & Trewhella 1988), which at the time considered both the use of vaccine and poison baits for foxes. It was also performed prior to the Animal By-Products (Enforcement) (England) Regulations 2011, which now prohibits putting animal by-products onto land where farmed animals could have access. However, improving bait uptake by foxes could also benefit other disease control scenarios, for example the distribution of anthelmintic baits for the control of *Echinococcus multilocularis* (Budgey, Learmount & Smith 2017), or indeed of improving bait uptake by other species; e.g. oral delivery of *M. bovis* vaccine in badgers (Palphramand *et al.* 2012).

Difference in the uptake of bait treated with both attractants or flavours, was compared with untreated bait in a rural area. The seasonal differences in the effectiveness of these additives in improving bait uptake by foxes was investigated by carrying out the experiments at 3 monthly intervals over a 15 month period.

## METHODS

### Attractants and flavours

These were identical to those used by Saunders and Harris (2000). Synthetic fermented egg (SFE) was prepared according to the recipe given in Turkowski *et al.* (1983). Valeric acid (VA) was bought from a chemical supply house (Sigma); beef flavour (Blend L202H) and honey flavour (D306K) was supplied by Master Taste, Dursley; UK.

### Bait preparation

Baits similar to those used by Trewhella *et al.* (1991) were prepared in advance and frozen. Untreated baits were prepared by mixing mechanically recovered chicken meat – MRM (Perrichicken - Perrimax Meat Company) with a 3% gelatin solution (300 bloom from porcine skin, Sigma) in the proportion of 4:1 (w/v) MRM to gelatin solution. Bait ingredients were mixed in bulk, dispensed into small food containers and frozen; each bait weighed approximately 62g (50g MRM plus 12ml gelatin solution). Attractant laced and flavoured baits were prepared in the same manner but the attractant or flavour was added to the gelatin solution before it was mixed with the MRM. SFE and VA baits contained approximately 0.5ml attractant per bait; beef and honey flavoured baits contained 0.05% flavour.

### Investigations into seasonal differences in bait uptake

The seasonal differences in the effect of attractants and flavours on bait uptake were tested by carrying out a replicate in each season. However, placing bait in the same study site at a 3 monthly interval may lead to increased bait uptake with time because of learning by resident foxes. Alternatively the use of a different site for each replicate may result in differences in bait uptake due to differences in fox numbers at the different sites. To allow for differences in bait uptake due to these factors, for each experiment a permanent site was chosen where replicates were carried out at 3 monthly intervals, and for each experiment for each season, a replicate was carried out at a single-use site; (in the flavour experiment it was not possible to carry out an autumn replicate at a single-use site). A further test for learning was done by repeating the first replicates in the next year.

### Study sites

Study sites were on farmland in Gloucestershire, England, and included both arable and grassland with small woodlands and uncultivated areas. The permanent study site for attractant baits was an arable farm and the field boundaries were a mixture of roadside grass verges, dry-stone walls and fences. The fields adjoined roads, woodland and pasture. The permanent site for flavoured baits was an ex-military airfield, part of which was used to grow crops and part was rough grassland. The airfield adjoined small woodlands and mixed farmland. The land used for the single use sites varied, some fields were permanent grass, whilst others were cropped and at various stages of growth or recently harvested.

### Bait placement procedure at study sites

At each site a circular route was chosen that followed field boundaries. Each route was divided into 15 sections with 5 bait stations per section. Within sections bait stations were spaced 100 paces apart with 200 paces between each section. Baits of the same type were used in each section; baits of different types were used in adjacent sections according to a rotating sequence (e.g. section 1 untreated, section 2 SFE, section 3 VA, section 4 untreated etc.) along the length of the bait line. At the permanent sites, in order to avoid placing the same type of bait in the same section at every replicate, the sequence was changed for each replicate; thus one type of bait was used on sections 1, 4, 7, 10, & 13 for one replicate but a different type of bait was used at those sections for the next replicate. Additionally, at the permanent sites, in order to prevent the repeated location of bait stations in exactly the same places; a slightly different start point was chosen each time a replicate was carried out. Thus although each section was in the same general part of the bait line throughout, from replicate to replicate the sections and bait stations were ‘staggered’.

On the first day of each replicate a single bait of the appropriate type was buried 10 cm deep at each bait station, following the old bait placement procedure for a rabies incident in the UK (Meldrum 1988). The location was marked by a number peg placed 1m from the bait. During each replicate, baits were inspected daily during daylight hours and missing baits replaced by the same type in the same hole. Baits were placed frozen as it was found that baits thawed within 2 hours of removal from the freezer. For each replicate, baits were placed and checked daily for 6-7 days. Thus for each bait type a total of 150-175 baits were available to be taken (6-7 days x 5 sections x 5 bait stations).

### Assessment of bait take

When bait was missing, a search for footprints or other field signs was made. Often there was little to indicate the species of animal responsible. The majority of baits were considered to have been removed by foxes because a mammal had obviously dug them up. However, the other species which could take baits (feral cats and badgers) would also be potential target species during a rabies baiting programme. At inspection the baits were scored as untouched, removed or present but disturbed. Additionally, at a few bait stations a camera was placed, which would be triggered when the bait was removed or disturbed, to obtain photographic records of animals removing buried bait. The field work was performed in the early 1990s.

## RESULTS

For the baits containing attractant, the permanent site showed a decrease in the total number of baits taken through the five seasons (Figure 1). This was true for untreated and treated baits. For the single-use sites there was a slight increase in the total bait take by season (Figure 1), and this trend was similar for all bait types. Therefore there was no evidence of learning. For the flavoured baits the permanent site showed an increase in the total bait take through the seasons (Figure 2), and this trend was seen for all bait types. The single-use sites showed a general decrease in bait take by season (Figure 2), which was also true for all bait types. It is therefore possible that the difference in bait uptake rates for the flavoured baits is due to learning, although this must have applied to all bait types.

**Figure 1.**
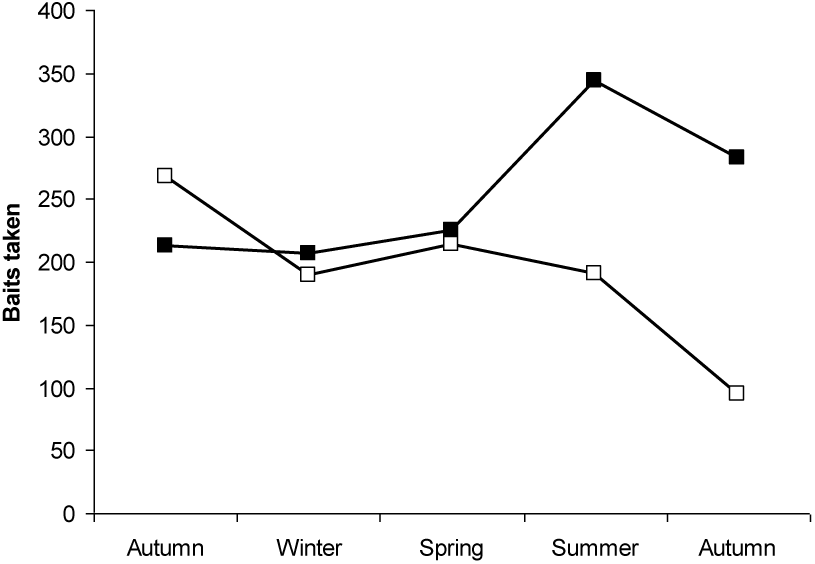
Total bait uptake in each season for the permanent site (open boxes) and single-use sites (closed boxes) for all attractant bait types (untreated, SFE and VA).

**Figure 2.**
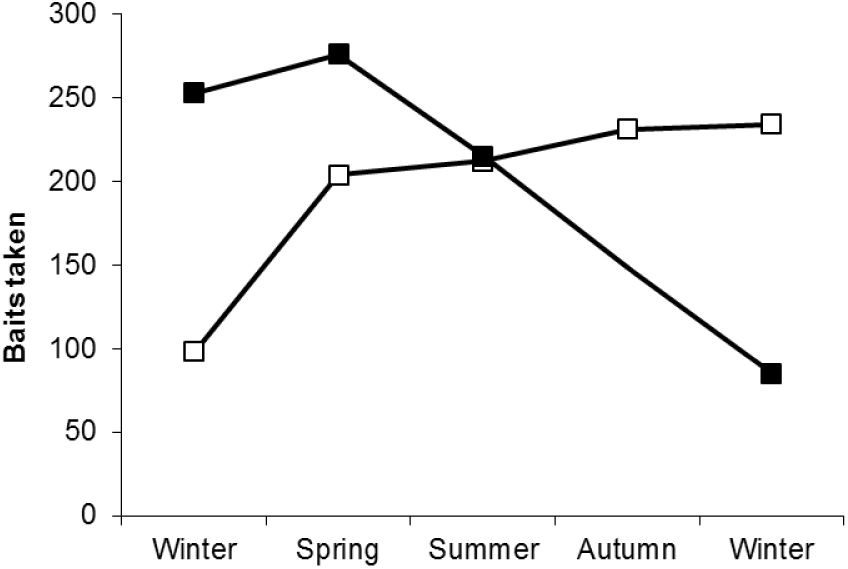
Total bait uptake in each season for the permanent site (open boxes) and single-use sites (closed boxes) for all flavoured bait types (untreated, beef and honey).

If we combine the data from all seasons and examine the increase in daily bait take for the first six days of baiting we expected to see an increase which may reach an asymptote by day six. For attractants at the permanent site this was clearly visible and occurred for all bait types (Figure 3). In this experiment the SFE baits were preferred over the other bait types. At the single-use sites no asymptote was apparent, but the SFE baits were clearly preferred over the other bait types (Figure 4). This approach was then used for the flavoured baits (Figures 5 and 6), which showed a preference for the honey baits at the permanent site and slight preference for the untreated baits in the single-use sites. In neither case was an asymptote reached during the six days.

**Figure 3.**
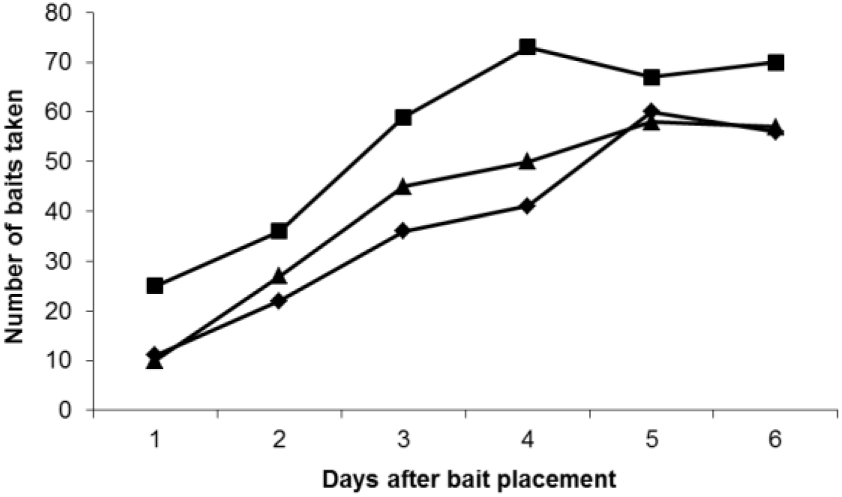
Daily total number of baits taken on each night at the permanent site for baits with added attractants (untreated: diamonds, SFE: squares, VA: triangles). Bait uptake is summed for the experiments in all five seasons.

**Figure 4.**
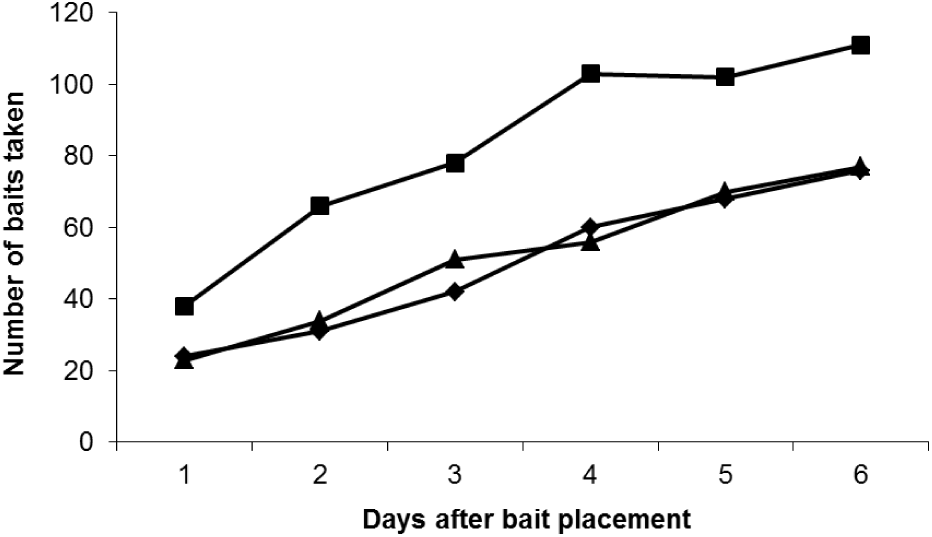
Daily total number of baits taken on each night at the single-use sites for baits with added attractants (untreated: diamonds, SFE: squares, VA: triangles). Bait uptake is summed for the experiments in all five seasons.

**Figure 5.**
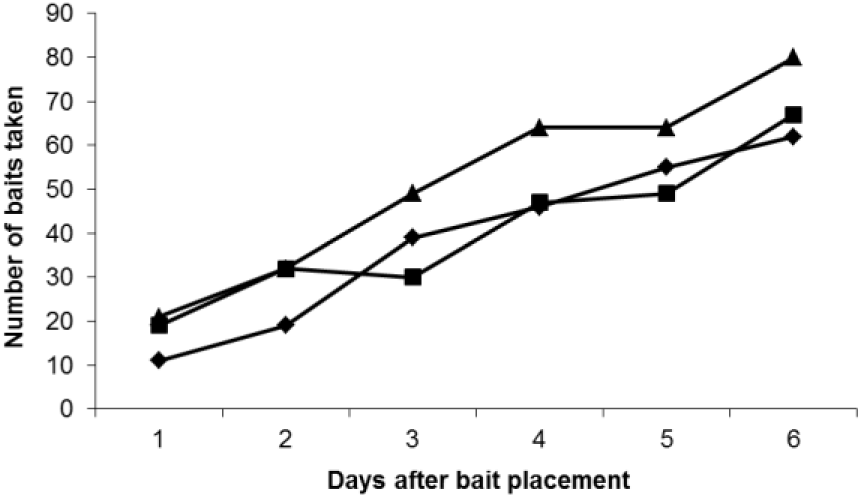
Daily total number of baits taken on each night at the permanent site for baits with added flavouring (untreated: diamonds, Beef: squares, Honey: triangles). Bait uptake is summed for the experiments in all five seasons.

**Figure 6.**
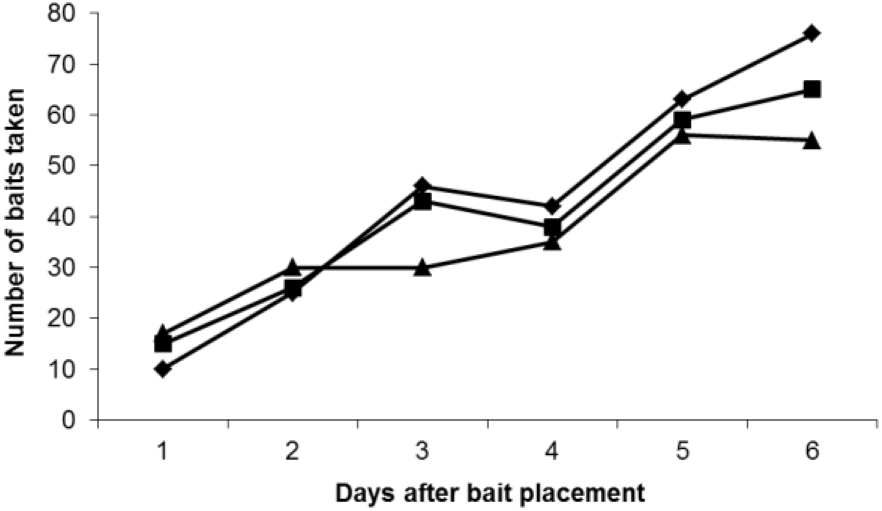
Daily total number of baits taken on each night at the single-use sites for baits with added flavouring (untreated: diamonds, Beef: squares, Honey: triangles). Bait uptake is summed for the experiments in all five seasons.

These results indicated some preference for the SFE-treated baits, but did not show that SFE-treated baits would lead to a higher kill rate if replaced by poison. For this we determined how often each bait type had the highest uptake rate on the last day of each of the 10 replicates for both experiments (five seasons at the permanent site and five seasons at the single-use sites: Table 1). A Monte Carlo randomisation test was then performed on the data. The first places (including ties) were attributed at random to each bait type 10,000 times and the probability that the experimental results occurred by chance were then exactly determined. The SFE baits were taken more often than expected by chance (p = 0.0474), and none of the other results were statistically significant.

**Table 1.**
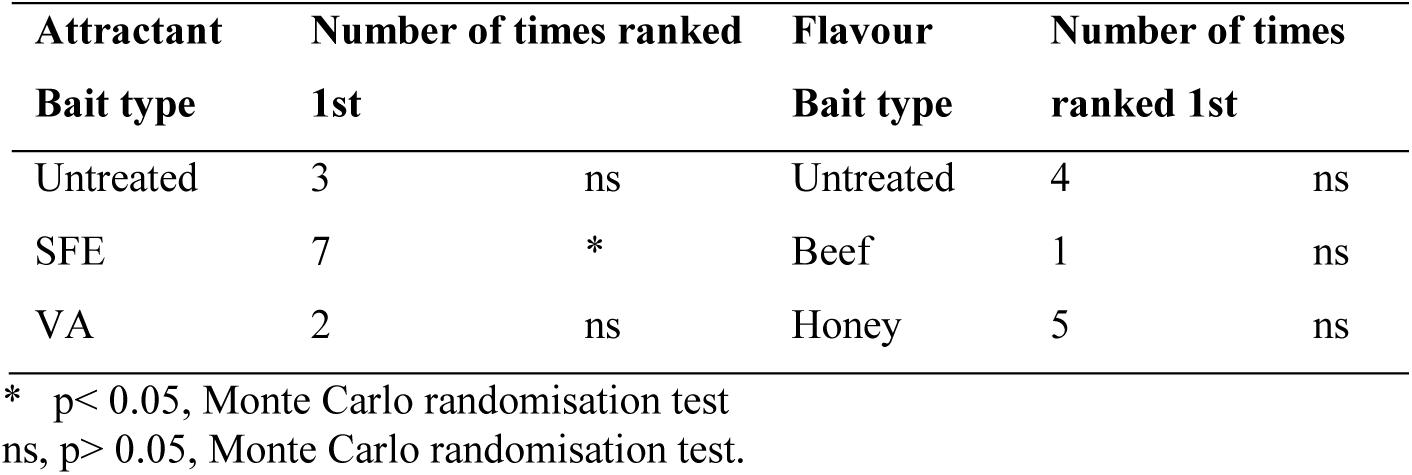
The frequency that each bait type had the highest uptake rate on the last night of the experiment. The attractant bait had ten experiments with two ties for first place and the flavoured bait had nine experiments with one tie for first place.

On the last day of baiting for the attractant-bait experiment a total of 132 untreated and 181 SFE-treated baits were taken. This represents a 37% increase in bait uptake on the last night. It should also be noted that more SFE baits were removed on the first night of the experiments than the other bait types (except for the summer replicate at the permanent site and the second autumn replicate at the single-use site). In the flavour-bait experiment, on the last day of baiting, almost equal numbers (138,132,135) of baits of any type were taken.

## DISCUSSION

### Species of animal removing bait

Very few field signs were found that indicated the species of animal responsible for removing the baits. At some bait stations considerable digging by animals attracted to the bait occurred; at others, especially where bait was bait buried under turf, the bait was removed with very little disturbance and it was necessary to lift the turf to determine whether or not the bait was still present. Both foxes and badgers are attracted to buried bait but the same animal does not always visit the same bait station on consecutive nights (Reynolds 2000). The camera recorded both foxes and badgers removing bait from the same bait station on different nights. Field signs following bait removal at these ‘camera traps’ were very similar irrespective of whether or not a fox or a badger removed the bait. The possibility exists that any footprints present at a bait station may be from the last animal to visit the bait station; thus a fox may remove a bait but subsequently a badger may be attracted to the bait station and leave its footprints. Consequently it must be concluded that field signs such as footprints are not very reliable indications of the species of animal removing the bait.

### Bait removal

At inspection, baits were recorded as present or removed (i.e. taken by foxes or other animals). Unlike the findings in urban areas, where partial consumption of bait was observed (Trewhella *et al.* 1991), on no occasions were partially eaten baits found. However, on a number of occasions baits had been disturbed but not removed; on these occasions the bait was found partially or totally exposed showing that it had at least elicited interest of an animal. No clear pattern to this phenomenon was apparent as it occurred at all sites with all bait types. There are a number of different possible reasons for this phenomenon. Chance disturbance of an animal, by agricultural or other human activities, taking bait may be the cause in some circumstances whilst in others satiated animals may continue to investigate food items.

### Frequency of bait removal from bait stations

Although individual baits were not removed until some days after first placement, when a bait was removed from a bait station it was frequently removed on subsequent days until the end of the period. Thus, in each replicate, a steady rise in the number of baits removed daily was observed; therefore in the short term foxes and other species quickly learn that food is available and exploit the situation, although the same animal does not always take bait from the same bait station on consecutive nights (Reynolds 2000). The frequency with which any animal visits a bait station is unknown. There may be inter- and intra-specific competition for bait. Some individuals may visit a bait station daily or more frequently but the first animal to arrive, following bait replenishment, is most probably the animal that removes the bait.

### Caching

It is well known that foxes will cache surplus food, including baits (Saunders, Kay & McLeod 1999; Thomson & Kok 2002), although there was no evidence whether caching occurred during this work. Pilot trials showed that raw chicken MRM was too fluid to be used as a bait on its own, especially during wet weather, and baits such as chicken heads tend to be carried away from the bait station to be eaten. This bait was deliberately designed to be friable, thus encouraging foxes to eat it *in situ* rather than carry it away, but robust enough to hold together during wet weather. However, direct observations of wild foxes eating gelatin stiffened MRM baits has shown that in some cases the bait was carried away; although the baits may have been eaten away from the bait station and not cached.

### Learning

In order to demonstrate that repeated baiting at one site leads to predator learning there would need to be an increase in bait uptake rates at the permanent sites in each experiment, when compared to the single-use sites. For the attractants this was clearly not the case, as the bait uptake reduced in each season at the permanent site. For the flavourings there was evidence of learning, as the bait uptake increased in each season, compared with a spring peak at the single-use sites. However, the evidence was not conclusive. The number of baits taken on the first night of baiting at the permanent site increased in each season, but the total bait uptake in the last season at the permanent site did not exceed the bait uptake in the first two seasons at the single-use sites (Figure 2). Therefore any learning by the local population of foxes was not strong, as the total bait take was within the variation of bait uptake seen at other sites. In order to be conclusive about animals learning to find baits we would also need to explain why learning occurred with flavourings but not attractants. Long-term learning may be site specific; some fox populations are subject to considerable control pressure, and long-term learning may be a factor at sites with stable populations, however many replicated experiments would be required to demonstrate this.

### Seasonality

Fox numbers naturally vary between study sites and seasons; the population is at its highest just after the breeding season and the greatest number of independent adult foxes in the late summer and early autumn make the greatest demand on food resources. Therefore the greatest uptake of bait may be expected during this period. Alternatively the greatest uptake of bait may be expected in the season that naturally occurring food is least abundant, usually assumed to be winter or early spring.

Considering total bait uptake (irrespective of bait type) over the 15-month period for each season for each of the two experiments (and considering the permanent sites separately from the single-use sites) bait uptake peaked in all four seasons. If the bait uptake from all experiments was combined there was a small peak in summer. Therefore we conclude that there was no consistent seasonal peak in bait uptake.

### Use of Additives to increase bait uptake

There was no clear preference for either of the flavoured baits, over the untreated baits, whereas there did appear to be a preference for the SFE-treated baits, which had a higher total number of baits removed, and was the most frequently removed bait on the last night. Other baiting trials have shown that each animal takes between 1.7 and 2.1 baits per night (unpublished data), so the additional 26 SFE baits that were removed could have been taken by between 12 and 15 animals. Alternatively, the 17% increase in bait uptake of SFE baits could account for an additional 8-10% of the population consuming baits (either vaccine or poison baits).

Unfortunately, this work was not able to measure the number of foxes taking bait, nor the number of foxes consuming baits that had been taken. There is clear evidence that foxes will eat preferred baits and cache less preferred baits when they are found (Saunders, Kay & McLeod 1999; van Polanen Petel, Marks & Morgan 2001). If the bait type taken most often is assumed to be the preferred bait, then the other bait types may be cached more often, thus increasing this difference between bait types. The compounds in SFE and VA are products of decay that would naturally occur in rotting meat baits. Thus it would be expected that foxes would be exposed to these compounds in naturally occurring food and therefore it can be expected that SFE and VA baits would be attractive to foxes.

Baits were placed where foxes tend to forage and travel but this alone will not ensure that foxes will find the baits. Detection of bait will be by smell; consequently it was possible that more SFE baits were taken not because they were preferred over other baits but because they were more easily found i.e. SFE baits could be detected from a greater distance than the other baits. This does not detract from the usefulness of an attractant like SFE for increasing bait uptake; baits other than SFE may be preferred or more palatable but if they are not easily found then their use will not improve bait uptake.

The main problem for control of a rabies outbreak in Britain was the low uptake of bait by foxes in urban areas (Trewhella *et al.* 1991) and not in rural areas, where these experiments were carried out. These experiments could not have been carried out at this scale, in an urban area without a considerable increase in resources, which was not available. The main aim of the work was to show whether or not use of any of the candidate additives could produce an increase in the uptake of treated bait over untreated bait. Subsequently experiments (similar to Trewhella *et al.* 1991) could then be carried out in urban areas using the additive that was shown to increase bait uptake by the greatest amount. The preferences for baits demonstrated here is likely to be valid in other habitats, and thus the addition of SFE to bait used in urban areas would be expected to significantly increase the proportion of the population consuming baits, although this has not been demonstrated.

SFE is an extremely unpleasant material to handle, having a persistent strong odour. It would only be possible to use SFE-treated bait if a method of producing treated bait in large quantities was developed. Such development work would be premature in the absence of evidence to show that the uptake of SFE treated bait, in urban areas is superior to that of any other bait. However, an additional advantage to the use of a compound like SFE would be that it is so unpleasant it would act as a deterrent to people; thus reducing the chance of human interference to bait with corresponding reduction to risk to human health especially if a poison bait were to be used.

## ACKNOWLEDGEMENTS

We would like to thank the various landowners for permission to carry out the work, and the Ministry of Agriculture, Fisheries and Food (now the Department for Environment, Food and Rural Affairs) for funding.

